# Analysis of convolutional neural networks reveals the computational properties essential for subcortical processing of facial expression

**DOI:** 10.1101/2023.01.18.524656

**Authors:** Chanseok Lim, Mikio Inagaki, Takashi Shinozaki, Ichiro Fujita

**Author notes:** **Corresponding author:** Ichiro Fujita, PhD, Laboratory for Cognitive Neuroscience, Graduate School of Frontier Biosciences, Osaka University, 1-4 Yamadaoka, Suita, Osaka 565-0871, Japan.

## Abstract

Perception of facial expression is crucial in the social life of primates. This visual information is processed along the ventral cortical pathway and the subcortical pathway. Processing of face information in the subcortical pathway is inaccurate, but the architectural and physiological properties that are responsible remain unclear. We analyzed the performance of convolutional neural networks incorporating three prominent properties of this pathway: a shallow layer architecture, concentric receptive fields at the first processing stage, and a greater degree of spatial pooling. The neural networks designed in this way could be trained to classify seven facial expressions with a correct rate of 51% (chance level, 14%). This modest performance was gradually improved by replacing the three properties, one-by-one, two at a time, or all three simultaneously, with the corresponding features in the cortical pathway. Some processing units in the final layer were sensitive to spatial frequencies (SFs) in the retina-based coordinate, whereas others were sensitive to object-based SFs, similar to neurons in the amygdala. Replacement of any one of these properties affected the SF coordinate of units. All three properties constrain the accuracy of facial expression information in the subcortical pathway, and are essential for determining the coordinate of SF representation.

## Introduction

Perceiving the facial expressions of other individuals plays a critical role in the social life of primates, including humans. Two neural pathways, the ventral cortical pathway and the subcortical pathway, contribute to this perceptual ability (Fig. 1A; Pessoa and Adolphs, 2010; Tamietto and de Gelder, 2010; Petray and Bickford, 2019). The ventral cortical pathway consists of a network of areas in the occipito-temporal region of the cerebral cortex, and processes a variety of visual features of objects, people, and environments, including shape, color, texture, material properties, and binocular disparity (Ungerleider and Mishkin, 1982; Connor et al., 2007; Conway et al., 2010; Roe et al., 2012; Kravitz et al., 2013; Vaziri et al., 2014; Verhoef et al., 2016; Komatsu and Goda, 2018). Neurons that preferentially respond to images of faces or facial features are found in several clusters along this pathway (Desimone et al., 1984; Perrett et al., 1987; Fujita et al., 1992; Haxby et al., 2000; Tsao and Livingstone, 2008; Duchaine and Yovel, 2015; Freiwald et al., 2016). They constitute the neural system that analyzes facial details such as expression, identity, and direction of attention. The subcortical pathway consists of a few processing stages in phylogenetically ancient regions: the superior colliculus in the midbrain, the pulvinar nucleus in the posterior thalamus, and the amygdala in the medial limbic system. The subcortical pathway is suggested to mediate rapid behavioral and physiological (autonomic) responses to sensory signals related to possible dangers such as fearful faces (Tamietto and de Gelder, 2010; Nakano et al., 2013; for a critical review, see Pessoa and Adolphs, 2010). The ventral cortical pathway and the subcortical pathway intersect at the amygdala.

**Figure 1.**
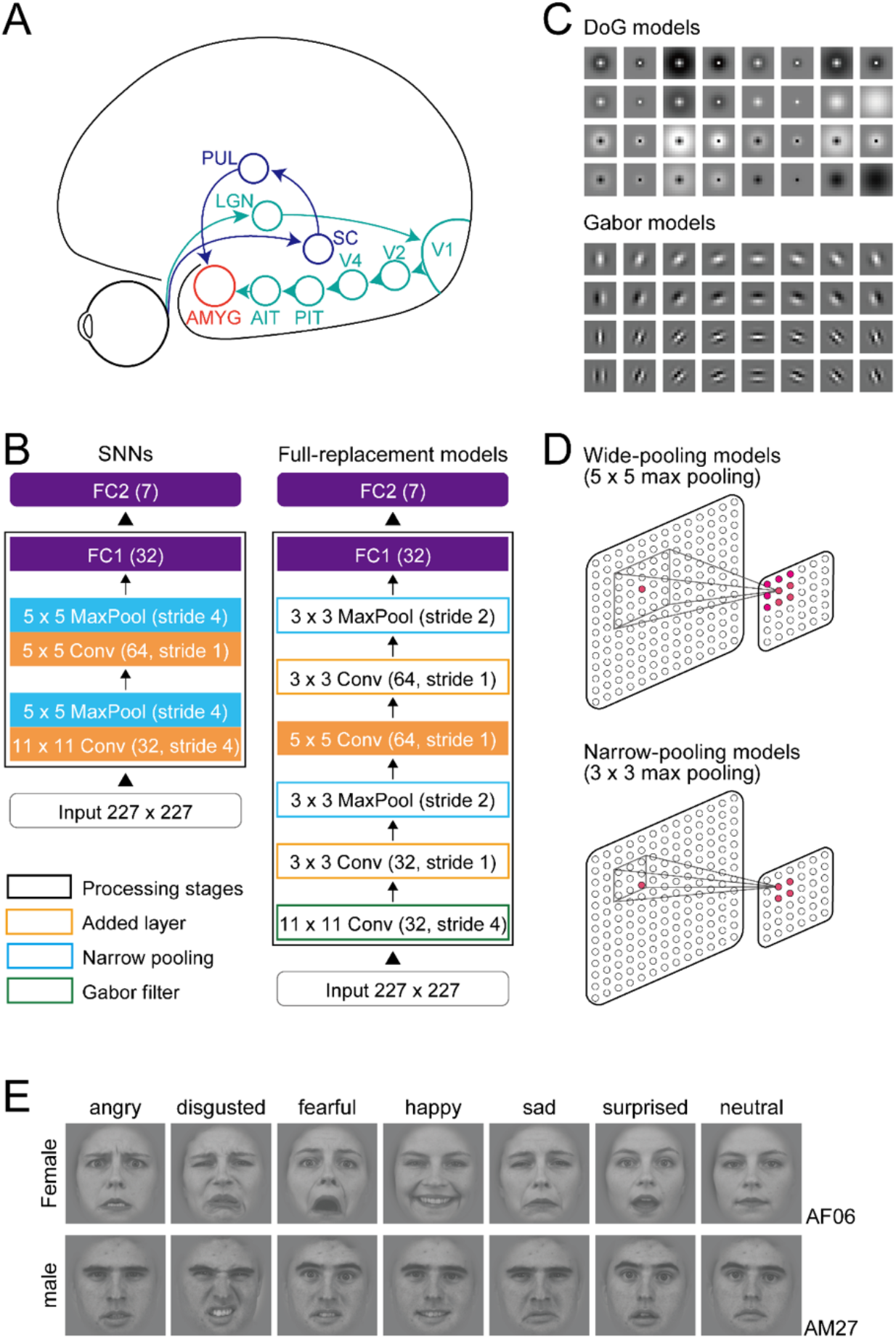
Shallow neural networks (SNNs) and modifications. (A) Cortical and subcortical visual pathways for processing facial expressions in the primate brain. AMYG: amygdala; AIT, PIT: anterior and posterior parts of inferior temporal cortex; LGN: lateral geniculate nucleus; PU: pulvinar nucleus; SC: superior colliculus; V1, V2, V4: visual areas 1, 2, 4. (B) A schematic illustration of the SNNs and full-replacement models. In the full-replacement models, processing layers were added, the filters in the initial layer were changed with Gabor filters, and the range of pooling was narrowed. (C) DoG filters for the SNNs and DoG models (upper) and Gabor filters for the Gabor models (lower). (D) The pooling range for the SNNs (5 × 5) and the narrow-pooling models (3 × 3). (E) Examples of presented face images with seven expressions (angry, disgusted, fearful, happy, sad, surprised, neutral) of two individuals (upper: female, AF06; lower: male, AM27). The original images were obtained from the Karolinska Directed Emotional Faces database (Lundqvist et al., 1998).

Psychological and brain imaging studies suggest that the subcortical pathway subserves the ability of some patients with lesions in the primary visual cortex (V1) to discriminate facial expressions despite lacking visual awareness (“affective blindsight”; deGelder et al., 1999; Pegna et al., 2005; Striemer et al., 2019). These patients also reflexively exhibit specific facial expressions and pupillary reactions when exposed to fearful or happy faces (Tamietto et al., 2009). Studies have also shown that the subcortical pathway supports unconscious face perception in neurologically healthy subjects (Morris et al., 1999, 2001). Furthermore, orientation bias toward faces or face-like patterns by newborn babies is suggested to be mediated by the subcortical pathway (Cassia et al., 2001; Johnson, 2005; but see Buiatti et al., 2019). Importantly, these perceptual abilities are not perfectly accurate, instead resulting in modest performance at above-chance levels. These findings suggest that information on faces conveyed by the subcortical pathway is less accurate than that carried by the ventral cortical pathway.

Electrophysiological studies have demonstrated that processing of facial expression in the subcortical pathway is indeed fast and not very accurate. Méndez-Bértolo and colleagues (2016) showed that intracranial local field potentials in the human amygdala respond differentially to fearful faces versus other faces within 74 ms after stimulus onset. A recent single-neuron recording study in the monkey revealed that a population of amygdala neurons responded to threatening faces even within 50 ms (Inagaki et al., 2022a). This early response, when combined across an ensemble of neurons, carries information that allows linear classifiers to discriminate threatening faces from neutral and affiliative faces. The rate of correctly discriminating the three expressions is around 50%; this is well above chance (33%), but significantly worse than perfect. Some neurons in the superior colliculus and pulvinar of the monkey also respond to faces and face-like patterns with an even shorter latency of 30–50 ms (Nguyen et al., 2013, 2014).

What architectural and physiological properties of the subcortical pathway are responsible for its fast, crude processing? The fast processing most likely arises from the small number, or “shallowness,” of processing stages in the subcortical pathway, given that the ventral cortical pathway and its upstream area (the lateral geniculate nucleus) consist of a larger number of regions (at least six before reaching the amygdala) than the subcortical pathway (only two), and that every transition from one cortical region to the next takes at least 10 ms (Schmolesky et al., 1998). It is unclear whether the shallow processing similarly explains the low accuracy of the information transmitted by the subcortical pathway to the amygdala. This uncertainty arises from the fact that in addition to the difference in the number of processing stages, visual response properties differ markedly between the two pathways. Neurons in the superior colliculus at the first stage of the subcortical pathway show circular receptive fields with center-surround organization, which can be modeled using the difference-of-Gaussian (DoG) function (Cynader and Berman 1972; Updyke 1974; Marino et al. 2008; Churan et al., 2012). By contrast, simple cells of V1 at the first stage of the cortical pathway have elongated receptive fields with side-by-side ON and OFF subregions, which can be modeled by two-dimensional Gabor functions (Jones and Palmer, 1987). Furthermore, the receptive field is typically larger in the superior colliculus (for the foveal field, 1.5–10° in superficial layers, 10–20° in deep layers; Goldberg and Wurtz, 1972; Wallace et al., 1997) than in V1 (1.18° in simple cells, 1.3° in complex cells; Van den Bergh et al., 2010) and the extrastriate areas V2 and V4 (Freeman and Simoncelli, 2011). Thus, spatial pooling across ascending stages occurs over a wider visual field area in the subcortical pathway than in the ventral cortical pathway.

In the present study, we addressed how these properties of the subcortical pathway, i.e., the shallowness of processing stages, DoG-type receptive fields at the initial stage, and spatial pooling over a wider visual field, influence facial expression processing. To this aim, we constructed convolutional neural networks (CNNs) and analyzed their performance in facial expression discrimination. CNNs are one type of multilayer perceptron, and can be optimized (“learn”) to classify inputs by varying connection weights between processing units through supervised learning algorithms (LeCun et al., 2015). Typical CNNs have several to tens of layers (deep neural networks, DNNs). DNNs developed for classifying visual objects share architectural and representational features similar to those of the ventral cortical pathway (Yamins et al., 2014; Güçlü and van Gerven, 2015; Yamins and DiCarlo, 2016; Hassabis et al., 2017). We designed our CNNs to imitate the subcortical pathway by reducing the number of processing stages and by implementing DoG-type receptive fields and a wider extent of pooling. These CNNs, hereafter referred to as shallow neural networks (SNNs), learned to discriminate facial expressions with modest correct rates. Replacing the three properties, one-by-one, two at a time, or all three simultaneously, with the corresponding properties in the ventral cortical pathway gradually improved discrimination performance, suggesting that all three features are responsible for limiting the performance of the SNNs. We further showed that like some neurons in the amygdala, a major group of units in the final processing layer of the SNNs were sensitive to spatial frequency (SF) in the retina-based reference frame as initially detected in the first processing layer, and that the three subcortical properties contribute to preserving the retina-based SF sensitivity.

## Materials and Methods

### Architecture of SNNs

We constructed SNNs incorporating the distinct properties of the primate subcortical pathway (Fig. 1B–D; Table 1). Unlike typical DNNs, the SNNs consisted of only two sets of convolution and pooling layers followed by two fully connected layers (FC1, FC2), approximating the small number of processing stages of the subcortical pathway. The first convolution layer incorporated 32 DoG-type filters (Fig. 1C, top) with a spatial resolution of 11 × 11 pixels, whereas weights in the second convolution layer were initially random, i.e., the filters had no structure, and gradually changed through training. A rectified linear unit (ReLU) was used as the activation function of a unit in the convolution layers and FC1; the ReLU forwards the processing results directly to the next stage if they are positive, otherwise it outputs zero. A max pooling operation was performed over 5 × 5 sliding regions with a stride of 4 for the outputs of convolution layers (Fig. 1D, top). Max pooling selected the largest value among the responses of units within a sliding window over the preceding convolution layer, and forwarded the value to the next layer. A local response normalization process was added after the pooling layers to aid generalization (Krizhevsky et al., 2012; we used slightly different parameters from theirs; k = 1, n = 5, *α* = 2 × 10^−5^, *β* = 0.75). Every unit in FC1 and FC2 received inputs from all units in the immediately preceding layer, i.e., each was fully connected. FC1 is the final processing layer, and FC2 outputs the results of entire processing by the SNNs. These features were implemented to capture the architectural and computational properties of the subcortical pathway, i.e., fewer processing stages compared to the ventral cortical pathway (Fig. 1A), DoG-type receptive fields in the superficial layer of the superior colliculus (Churan et al., 2012), and large receptive fields of deeper superior colliculus neurons (Wallace et al., 1997). The first three processing layers were intended to represent the superior colliculus, pulvinar, and amygdala, respectively. The processing types of these layers, i.e., convolution and pooling in the first two layers and full connection in FC1, were chosen to match the retinotopic organization of the three brain regions. The convolution and pooling processes in the first two layers exploit retinotopy, as the superior colliculus and pulvinar contain retinotopic maps (Bender, 1981; Chen et al., 2019). The FC1 layer loses retinotopic information because of the fully convergent connection from the earlier stage, as the amygdala does not have a retinotopic map (Morawetz et al., 2010).

**Table 1.** Architecture of the Shallow Neural Network (SNN). Each row describes a layer *i* with calculation operator Fi, output resolution *H*_*i*_ × *W*_*i*_, and the number of output channels *C*_*i*_. Conv denotes convolution layer, and Pool denotes max pooling layer.

| Input &<br>Layer<br>$i$ | Operator<br>$F_i$ | Resolution<br>$H_i \times W_i$ | #Channels<br>$C_i$ |
| --- | --- | --- | --- |
| Input | | $227 \times 227$ | 3 |
| Layer 1 | $11 \times 11$ Conv (stride 4) & $5 \times 5$ Pool (stride 4) | $55 \times 55$ | 32 |
| Layer 2 | $5 \times 5$ Conv (stride 1) & $5 \times 5$ Pool (stride 4) | $4 \times 4$ | 64 |
| Layer 3 | Fully connected | $1 \times 1$ | 32 |
| Layer 4 | Fully connected | $1 \times 1$ | 7 |

The SNNs were trained to discriminate images of facial expressions representing seven basic emotions: angry, disgusted, fearful, happy, sad, surprised, and neutral (Fig. 1E; see below for details). For each input image, the seven units in FC2 yielded scores ranging from 0 to 1 for the seven expression categories, representing the probabilities of classified expressions. The expression with the highest score was taken as the output of the model.

The DoG-type filters of the first convolution layer were built using the following formula:

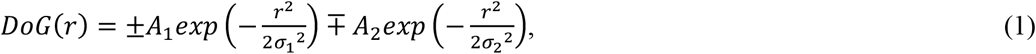

where *r* is the polar radius from the filter center, *A*_*1*_ and *A*_*2*_ are the amplitudes of exponentials of two Gaussian functions, and *σ*_*1*_ and *σ*_*2*_ are the standard deviations. Values of *A*_*1*_, *A*_*2*_, *σ*_*1*_, and *σ*_*2*_ were chosen empirically so that DoG curves took the shapes of Mexican hats. *A*_*1*_ values were 0.4, 0.67, 0.8, and 1.0. *A*_*2*_ values were determined based on *A*_*1*_ *− A*_*2*_ = 0.4. When *A*_*1*_ was 0.4 (i.e., *A*_*2*_ is 0, and *σ*_*2*_ cannot be defined), we set the *σ*_*1*_ value at 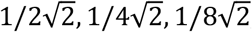, or 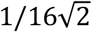. Otherwise, the *σ*_*1*_ value was 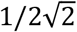 or 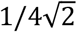. The *σ*_*2*_ value was based on *σ*_*1*_ */ σ*_*2*_ *=* 0.5 or 0.25. The same number of filters was generated for each *A*_*1*_ value.

We also constructed modified models in which the three properties of the SNNs were replaced one-by-one, two at a time, or all three simultaneously with the corresponding properties in the ventral cortical pathway. First, we added additional convolution layers with filters of 3 × 3 pixels after each of the first two convolution layers to increase the number of processing stages (add-layer model). In adding the convolution layers, the stride of sliding filters was reduced to 1 to keep the output resolutions unchanged before and after adding the new layers. Also, to keep the number of the output channels unchanged, the new layers contained the same number of filters as the preceding layers.

Second, we replaced the DoG-type filters with Gabor-type filters (Gabor model). Gabor-type filtering occurs in simple cells of V1, and emerges in the first layer of DNNs after they are trained to classify object images (Krizhevsky et al., 2012; Rai and Rivas, 2022). We constructed the Gabor-type filters with the following formula:

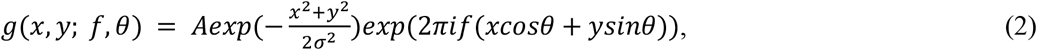

where *A* is the amplitude of the Gaussian envelope, *i* is √*−*1, *f* is the carrier frequency of a Gabor filter, and *θ* is the orientation (Movellan, 2002). *A* was fixed at 0.4 to match the amplitude of the DoG filters. *σ* was fixed at 0.125 so that the half-amplitude width was half of the filter width. *f* values were 2 or 4 cycles/image. Orientation *θ* was 0, 22.5, 45, 67.5, 90, 112.5, 135, or 157.5. We built even-and odd-symmetric filters for every combination of variables. In total, we obtained 32 Gabor-type filters (Fig. 1C, bottom).

Finally, we made the pooling window (convergence field) in the max pooling layers smaller (3 × 3; Fig. 1D, bottom) than that of the SNNs (5 × 5; Fig. 1D, top), enabling better spatial resolution of processing (narrow-pooling model) to mimic the smaller receptive fields in the visual cortices compared to the superior colliculus and pulvinar (Wallace et al., 1997; Vand den Bergh et al., 2010; Freeman and Simoncelli, 2011). The pooling range of 3 × 3 is often used in DNNs (e.g., AlexNet of Krizhevsky et al., 2012; ResNet of He et al., 2016).

### Face images and training of the SNNs

Face images were obtained from Karolinska Directed Emotional Faces (KDEF; Lundqvist et al., 1998) and Radboud Faces Database (RaFD; Langer et al., 2010). Images of the seven expressions of 40 individuals (half females, half males) were chosen from each database (the total number of images was 560 = 7 × 40 × 2). We converted the images from color into grayscale, and extracted the face region by removing hair, neck, and ears with the face-detection function of a computer vision library, OpenCV (Open Source Computer Vision Library; Bradski, 2000). The isolated faces were pasted on a gray background (198 × 198 pixels; RGB values = 128; Fig. 1E). We augmented the number of face images by changing size and position, and by flipping horizontally; seven sizes (28 × 28, 56 × 56, 85 × 85, 113 × 113, 141 × 141, 170 × 170, and 198 × 198 pixels), five positions (center, left-top, right-top, left-bottom, and right-bottom; directional displacements = 10 pixels), and two horizontally flipped images. The augmentation increased the number of images by 70 times to 39,200. At each training session, we randomly split this augmented set of face images into a training set (29,400 images), a validation set (2,450 images), and a test set (7,350 images). The number of images per facial expression was identical within each of these stimulus sets. To avoid the inadvertently biased assignment of face images of a particular size, position, or horizontal flip state into a given set, all images from the same individual were assigned into the same set.

The training was performed through supervised learning, and was conducted individually 20 times with randomized initial weights except for the built-in weights of the fist convolution layer, i.e., 20 SNNs with different initial states were built. In training, the weights other than the first convolution layer were optimized for classification of face images into the seven categories. Stochastic gradient descent was used for weight optimization. For each iteration, 32 samples were randomly selected from the training set as a mini-batch. The averaged cross-entropy (Goodfellow et al., 2016) across the 32 images in a mini-batch was calculated as an estimate of loss value, which is a measure of the difference between a model output and a supervised signal, and is used for quantifying the training effect. The number of iterations (i.e., weight-updating processes with single mini-batches) was set at 240,000. Initial weight parameters followed a normal distribution with a mean of 0 and a standard deviation of √(*2/N*) (*N* is the total number of weights; He et al., 2015). Weights were updated at each iteration with a constant learning rate of 0.001. This learning rate was determined empirically; a preliminary analysis based on 10 constructed SNNs (different from the 20 SNNs in the main analysis) revealed no decrease in loss values (i.e., no learning) with a learning rate of 0.01, which has frequently been used for DNNs in the literature (e.g., Simonyan and Zisserman, 2014). A dropout process was added before FC2 to facilitate learning across all units. The proportion of units dropped out of each weight update was set to 0.5. The training was conducted in a Python environment (Chainer 3.0.0; Tokui et al., 2015) on a graphics processing unit (GPU) machine (Intel^®^ Core™ i7-5820K Processor. Intel, Santa Clara, CA, USA; The GeForce^®^ GTX 1080 Ti, NVIDIA, Santa Clara, CA, USA). While the SNNs were being trained with the training set, the correct rate and loss value for the validation set were periodically checked to monitor signs of overfitting. After training was completed, the performance of the models was evaluated using the test dataset that had not been used for training. This was done to ensure that the models acquired a genuine ability to classify the facial expressions, as opposed to simply sorting the training images into the seven facial expression categories according to the instruction signals.

### Test for reference frames of SF tuning of model units

A difference in visual responses between the two pathways is the reference frame of neuronal tuning to SFs (Inagaki and Fujita, 2011). Neurons in the inferior temporal cortex, the final stage of the ventral cortical pathway, are tuned to object-based SFs (cycles/object) and represent face patterns in a size-invariant, hence distance-invariant, manner (Fig. 2A, right). Thus, the ventral cortical pathway converts the representation of SFs in the retina-based coordinate (cycles/degree) to that of object-based SFs. By contrast, many amygdala neurons preserve sensitivity to retina-based SFs. When the stimulus size is changed, these neurons change their preferred object-based SFs; for large stimuli, they respond to higher object-based SFs, which correspond to the same retina-based SFs (Fig. 2A, left). We analyzed the reference frame of FC1 SF tuning to evaluate how well our models captured this characteristic of subcortical processing.

**Figure 2.**
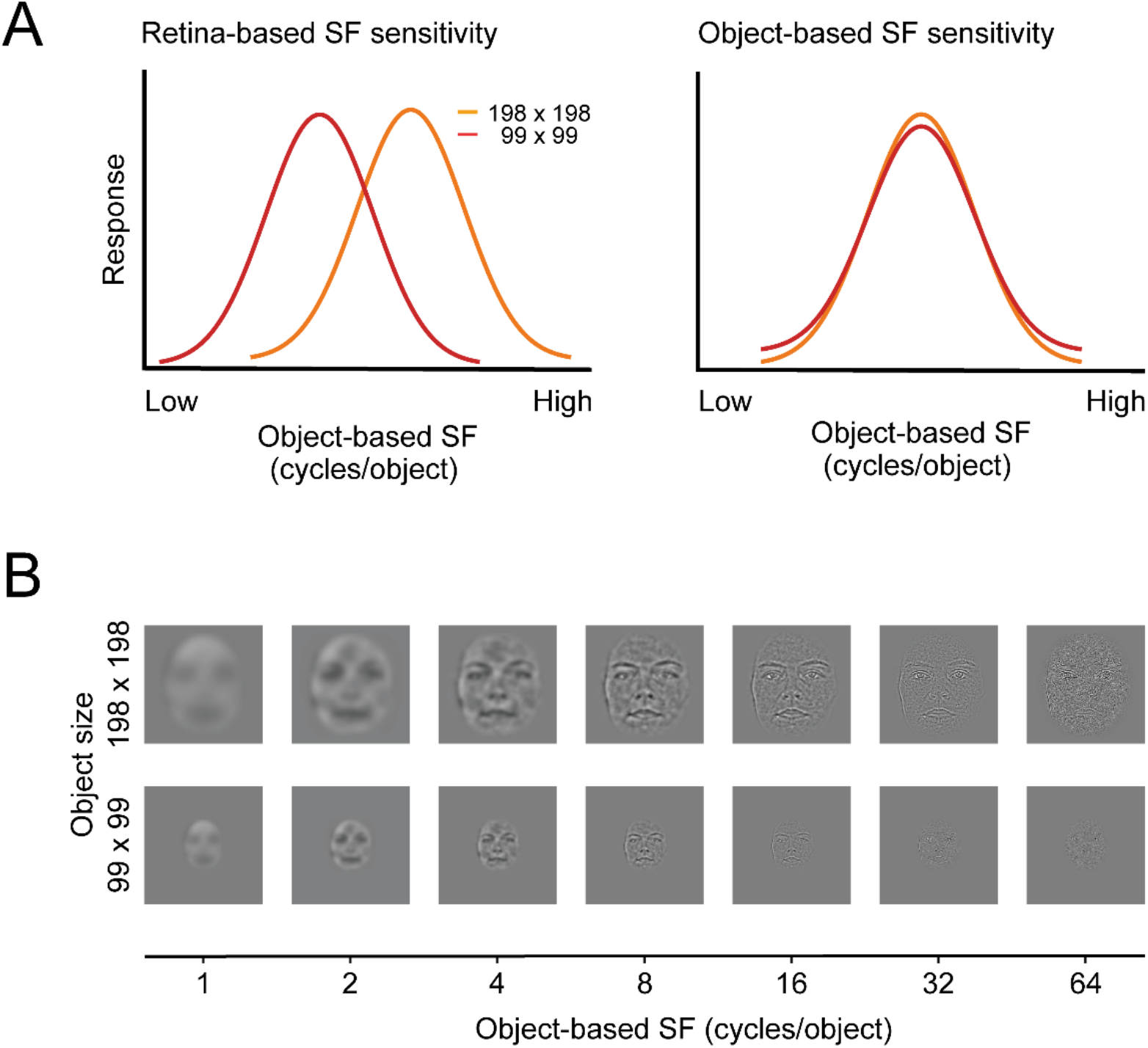
Test determining whether units are tuned for object- or retina-based spatial frequencies (SFs). (A) Hypothetical tuning curves for object-based SFs of units ideally tuned to retina-based SFs (left: peak shift = 1) or object-based SFs (right: peak shift = 0). (B) The models were fed face images with two different sizes (198 × 198 and 99 × 99 pixels) and 64 different bandpass filtering. These images were created by applying two-dimensional bandpass filters that shared the same center object-based SF across different sizes. For each unit in FC1, we obtained responses to images of different center SFs to create tuning curves for object-based SFs.

Bandpass-filtered face images were used to examine the SF tunings of FC1 units (Fig. 2B). These images were created by multiplying Gaussian functions with the original face images on the polar Fourier domain. Gaussian functions had 61 different center frequencies between 1 cycle/object and 64 cycles/object. The center frequencies had discrete values at steps of 0.1 cycles/object on a log scale. Gaussian functions shared the same variance at 2.4 octaves, regardless of their center frequencies. The filtered images had amplitude spectra that were determined solely by the Gaussian function because their spectra were set to be flat before the multiplication. To balance the total luminance contrast among the filtered face images, the peak amplitude of the Gaussian function was set inversely proportional to the center image-based SF (Inagaki and Fujita, 2011). These bandpass-filtered images were created for the seven facial expressions at two different sizes (99 × 99 and 198 × 198 pixels).

To characterize the reference frame of SF tuning of each unit, the peak SFs for the two stimulus sizes were estimated, defined by the SFs at which filtered face stimuli activated a unit most strongly. For a given unit, 14 peak SFs were determined (two sizes × seven expressions). The degree to which unit responses to SFs depended on the stimulus size was quantified by calculating differences in peak SFs on a log scale between the two face sizes. A peak shift of 0 indicates that a unit respond to the same cycles/object regardless of the image size, and is perfectly tuned to object-based SFs (Fig. 2A, right). A peak shift of 1 means that an SF tuning curve shifts by the amount corresponding to the change in the stimulus size, indicating that a unit is perfectly tuned to retina-based SFs (Fig. 2A, left). This analysis excluded cases in which units did not respond to face images or were not sensitive to SFs, and cases in which peak SFs were at either end of the tested range of SFs and the peak positions could not be determined.

### Analysis of the effects of the max pooling operation on SF selectivity

The max pooling operation collapses positional information of edges, which is detected by convolution filters and is critical for encoding the SFs of facial images. We therefore examined the effects of the max pooling on the SF selectivity of units. We were particularly interested in the role of the max pooling in converting sensitivity to retina-based SFs to sensitivity to object-based SFs. Bandpass-filtered images of the two stimulus sizes (198 × 198 and 99 × 99 pixels) were fed to the models, and it was determined how different stimulus sizes affected the responses of units to SFs in the first convolution layer (before pooling) and the first max pooling layer (after pooling). Changes in response patterns across the units associated with different stimulus sizes were quantified by calculating the dissimilarity index. The dissimilarity index *D* (*x, y*) for responses *x* to the large stimuli and responses *y* to the small stimuli was defined by the Euclidean distance between *x* and *y* as follows:

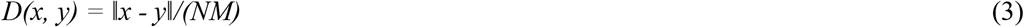

where ∥·∥ is the Euclidean distance, and *N* is the number of elements of *x* and *y. M* is the maximum value among the 6,405 Euclidean distances calculated for 61 center SFs and the seven facial expressions of 15 individuals. To probe the roles of the max pooling, the ratio of the dissimilarity index before pooling in the first convolution layer and after pooling in the first max pooling layer was then calculated. This analysis was applied to the case of wide pooling (5 × 5) and narrow pooling (3 × 3), as well as to the case of the SNNs and the Gabor models. By dividing ∥x - y∥ by *M*, the dissimilarity index was normalized across the layers (convolution vs. max pooling) and the models (SNNs vs. Gabor models), taking values from 0 to 1.

## Results

### Performance of SNNs in facial expression classification

The SNNs were trained to classify each face image in the training set into one of the seven facial expressions. The training improved the classification performance rapidly over the initial iterations and then slowly thereafter. The correct rate across the seven facial expressions rose from the chance level (0.14), surpassed 0.6 around 50,000 iterations, further improved to around 0.8 over 150,000 iterations, and reached an asymptote (Fig. 3A for an example SNN; 3B for the average of the 20 constructed SNNs; orange lines). The correct rate for the validation set saturated at around 0.5, which was substantially lower than for the training set, indicating insufficient generalization to “unseen” images. However, the validation correct rate reached a plateau in a similar way to the training correct rate. This indicates that the low correct rate was not the result of inadequate training, but represents the limited learning ability of the SNNs. It also indicates that no overfitting occurred. The loss value also quickly decreased over the initial 50,000 iterations, and became gradually stable (Fig. 3A, B; cyan lines). The results indicate that within the range of adopted iterations (240,000), the SNNs were trained to classify facial expressions without overfitting.

**Figure 3.**
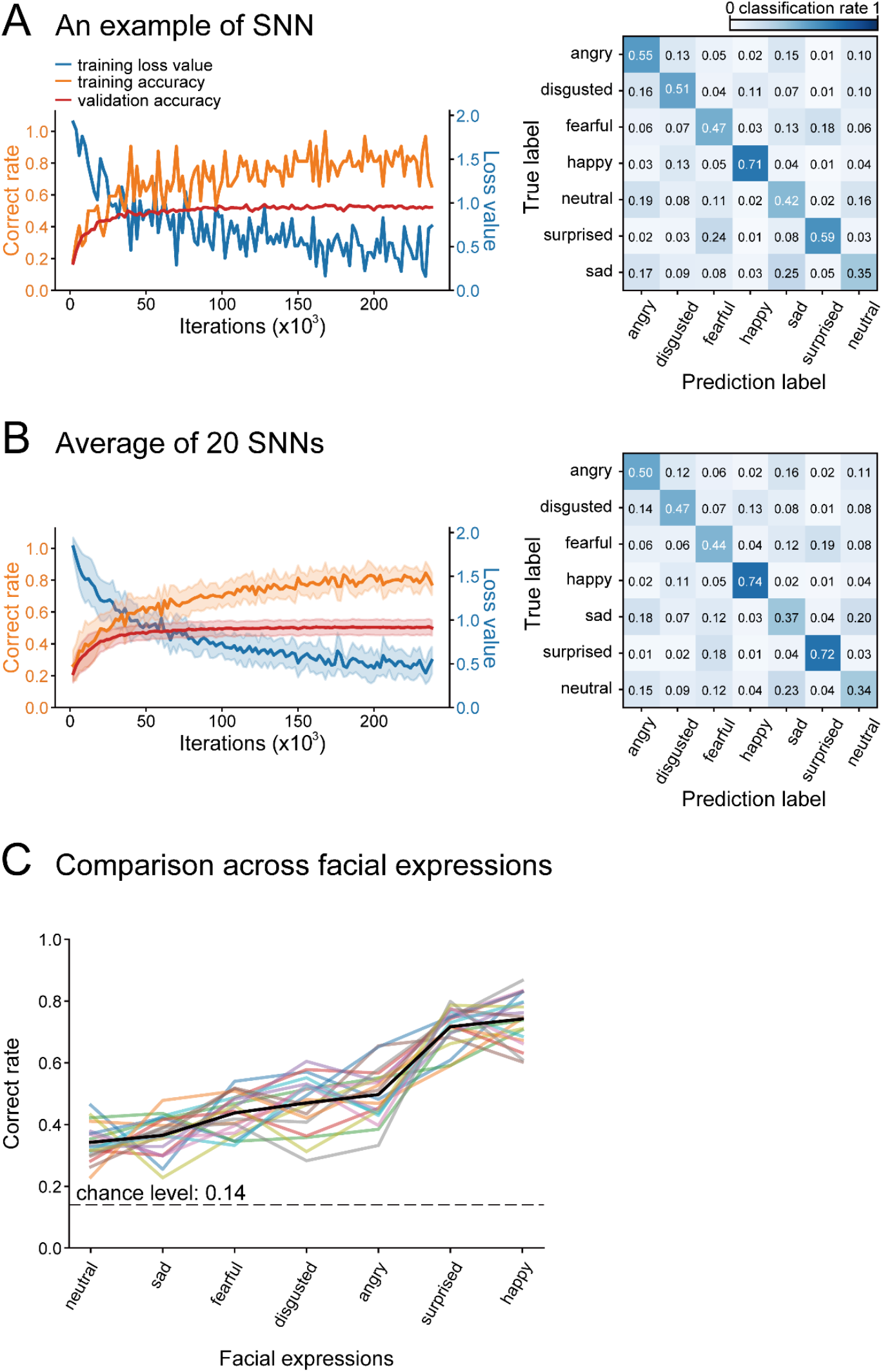
Learning curves and confusion matrices of an example SNN (A) and the average of 20 SNNs with different initial weights (B). Left panels show changes in correct rates that occurred during training in the training set (orange) and the validation set (red), and loss values (cyan). Confusion matrices on the right indicate the rate of classification of each facial expression (true label) as one of the seven expressions (prediction label). (C) The correct rates for the seven expressions. Black line indicates the mean of the 20 SNNs, and lines with other colors indicate the individual performance of the 20 SNNs. The order of facial expression is based on the mean correct rate.

The correct rates for the test set were higher than the chance level (1 / 7 = 0.14) for all facial expressions, but were modest; the average correct rate across the seven expressions was 0.51 for the example SNN shown in Fig. 3A. The average correct rate across the 20 constructed SNNs was 0.51 (± 0.03, s.d.). This was not different from the average correct rate across 20 additional SNNs that were trained with 3,000,000 iterations (0.50 ± 0.03; *p* = 0.289, *t*-test). The training performance thus did not improve even when the SNNs underwent overly excessive training, assuring that the modest correct rate was not due to insufficient training but instead reflected the limited ability of the SNNs. Confusion matrices showed that the correct rates varied among the facial expressions (Fig. 3A, B, right panels). Based on the averaged performance, the classification performance of the SNNs was best for happy (0.74) and surprised faces (0.72), followed by angry (0.50), disgusted (0.47), and fearful (0.44) faces, and was worst for sad (0.37) and neutral (0.34) faces. Sad faces were often confused with neutral, angry, and fearful faces. Neutral faces were often confused with sad and angry faces. This expression-dependent performance was consistent across the 20 constructed SNNs (Fig. 3C; *p* < 0.001 for expressions, *p* = 0.562 for models, two-way ANOVA).

### Effects of modification of SNNs on classification performance

Replacement of one or more of the three subcortical properties, namely the shallowness of processing stages, the DoG-type receptive fields at the initial stage, and spatial pooling over a wider visual field, with the corresponding cortical properties improved the classification accuracy (Fig. 4A; *p* < 0.001, ANOVA). The correct rates averaged across the seven expressions and the 20 constructed models of each modification were increased from 0.51 in the SNNs to 0.55 in the narrow-pooling models, 0.64 in the Gabor models, and 0.69 in the add-layer models (*p* < 0.01 / 28=_8_C_2_; *t*-test with Bonferroni correction). Among these models with one replaced property, the add-layer models exhibited the best performance. The results indicate that all three properties had effects on classification performance, and the layer structure was the most influential.

**Figure 4.**
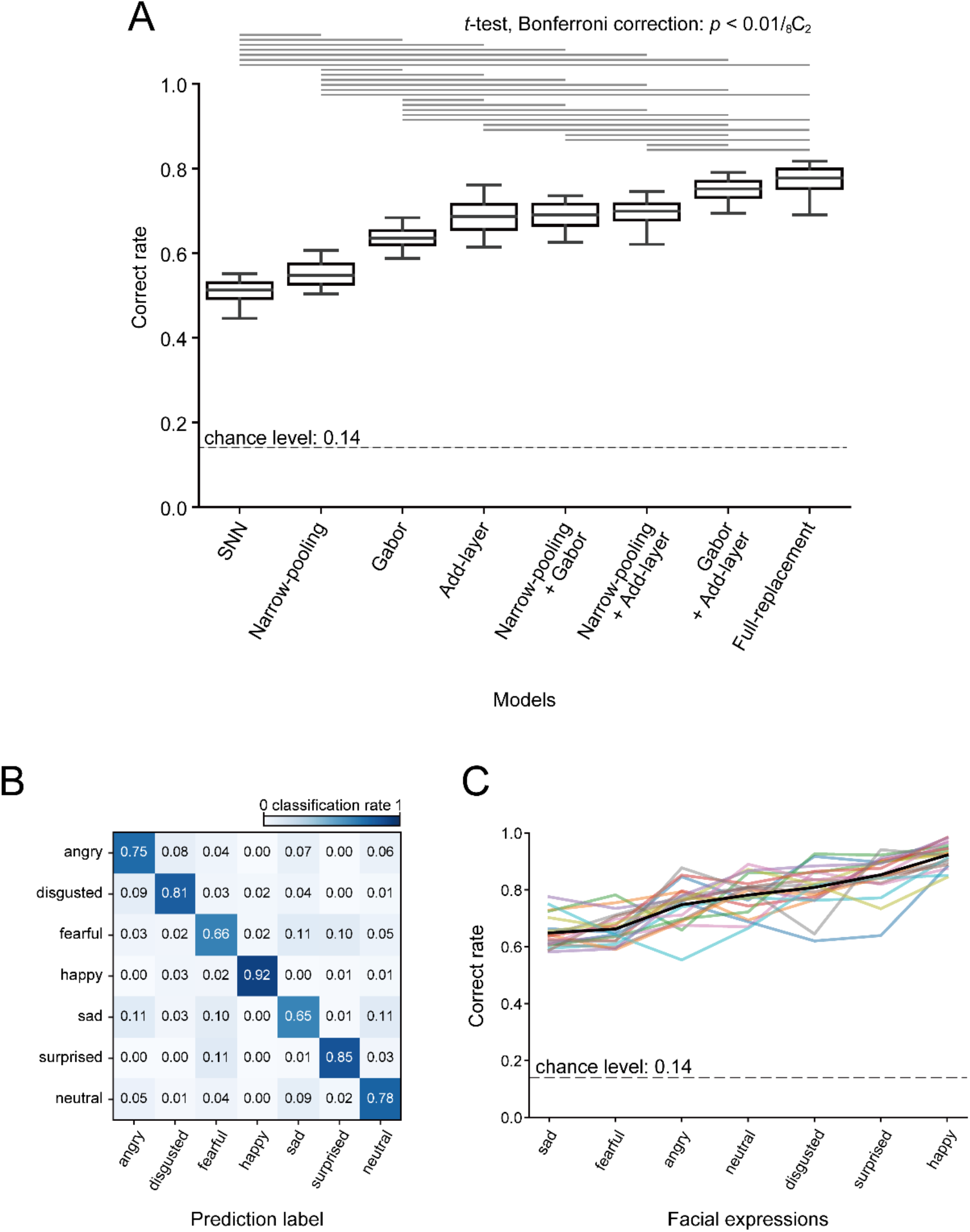
Effects on model performance of replacing subcortical properties with corresponding cortical properties. (A) Discrimination performances of the SNNs and the modified models. The discrimination performance differed across the models (ANOVA, *p* < 0.001). The pairs of models with statistically significant differences in the performances are linked with horizontal lines in the upper part (*t*-test, Bonferroni correction, *p* < 0.01/ 28=_8_C_2_). (B) Confusion matrix for the full-replacement models (average of the 20 constructed models). (C) The correct rate for the seven expressions across the 20 full-replacement models. The black line indicates the mean, and lines with other colors indicate the data for individual full-replacement models.

When two properties were replaced together, the narrow-pooling + Gabor models and the narrow-pooling + add-layer models performed better than the narrow-pooling models and the Gabor models (*p* < 0.01 / 28=_8_C_2_; *t*-test with Bonferroni correction), but comparably to the add-layer models (correct rate, 0.69, *p =* 0.778 for the narrow-pooling + Gabor models; 0.69, *p =* 0.564 for the narrow-pooling + add-layer models). The Gabor + add-layer models performed better than all one-property-replacement models (correct rate, 0.75; *p* < 0.01 / 28=_8_C_2_). When all three properties were replaced together (full-replacement models), the correct rate was 0.77, which was better than all other models (*p* < 0.01 / 28=_8_C_2_) except for the Gabor + add-layer models (*p* = 0.00846 > 0.01 / 28=_8_C_2_). The performance was improved for all facial expressions (Fig. 4B; mean correct rates: happy = 0.92; surprised = 0.85; disgusted = 0.81; neutral = 0.78; angry = 0.75; fearful = 0.66; sad = 0.65). As in the SNNs, it was highest for happy and surprised faces, and lowest for fearful and sad faces. The performance was improved most for neutral faces (SNNs, 0.37; full-replacement models, 0.78). The variance of the correct rates was affected both by facial expressions and models (Fig. 4C; *p* < 0.001 for facial expressions, *p* = 0.00285 for models, two-way ANOVA). The improved performance of the two-property–replacement and full-replacement models indicate that the effects of the three features on classification performance were partially additive, suggesting that the three features exerted their effects partially independently.

### Spatial frequency representation in the FC1 layer

As shown above, the SNNs exhibited modest performance in facial expression classification, and this performance was improved by changing SNN subcortical properties to corresponding cortical properties. These findings suggest that the SNNs captured aspects of processing in the subcortical pathway to the extent that they explained the suboptimal perceptual performance of V1-lesioned patients. We next looked into individual computational units to gain insights about the processing in the models. We examined SF sensitivities of FC1 units using two different sizes of input images (198 × 198 and 99 × 99 pixels). This procedure allowed us to determine whether units were sensitive to retina- or object-based SFs (Fig. 2; see Materials and Methods).

FC1 units of the SNNs exhibited a variety of dependencies of SF tunings on stimulus size (Fig. 5A). Some units responded to the same range of object-based SFs for both large and small stimuli, and the peak positions of the SF tuning curves remained unchanged (Fig. 5Aa, Ab). Other units exhibited different preferred SFs for large and small stimuli, and in these cases the peak position shifted horizontally along the abscissa (Fig. 5Ac–f). We quantified these shifts by measuring the difference between preferred SFs on a log scale for the two stimulus sizes. A peak shift of 0 means that the unit encoded SFs in the object-based coordinate, whereas a peak shift of 1 means that the unit encoded SFs in the retina-based coordinate. The peak shifts of the example units shown in Fig. 5A were 0.0 (a), 0.1 (b), 0.1 (c), 0.9 (d), 1.0 (e), and 1.3 (f).

**Figure 5.**
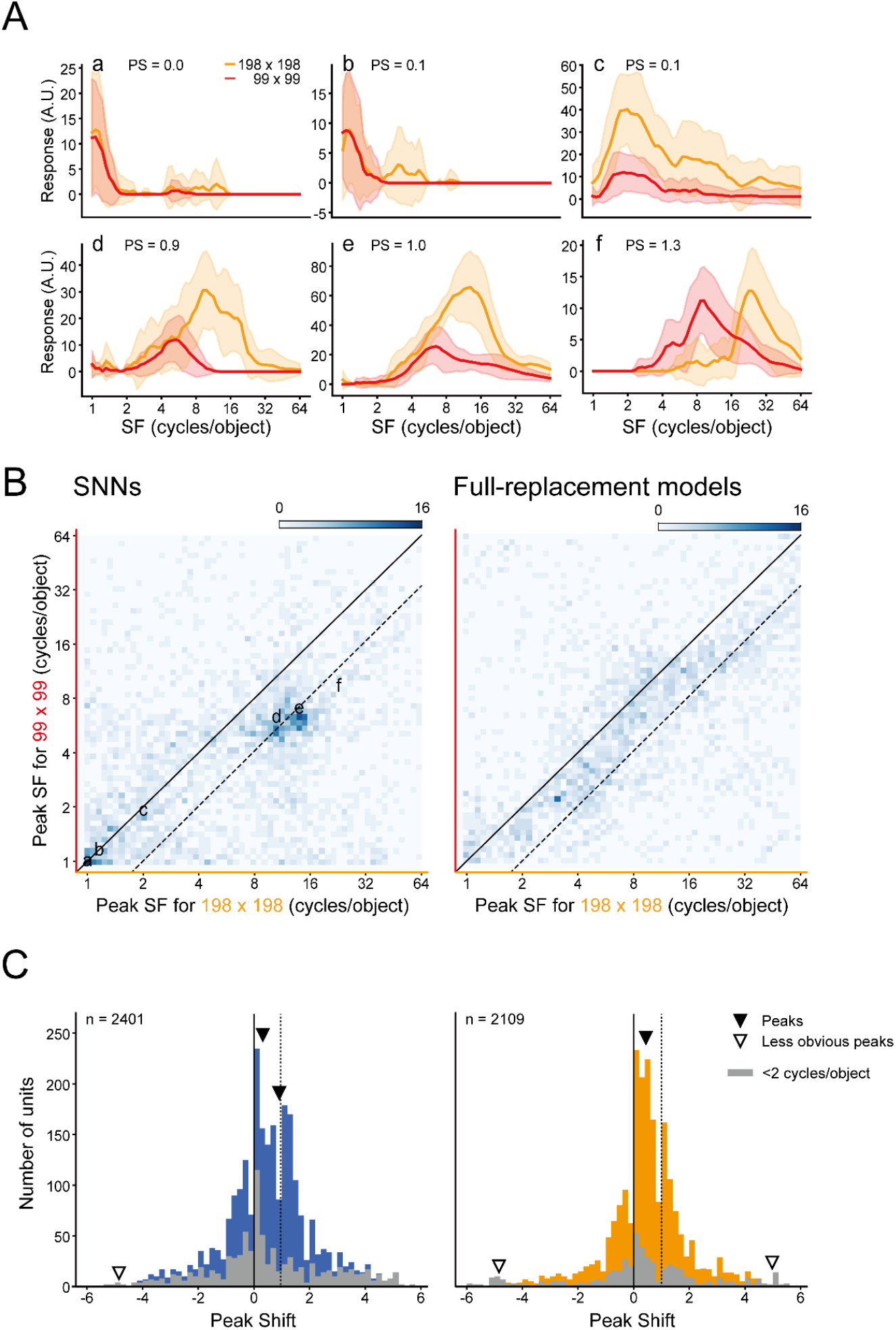
SF tuning reference frames of FC1 units of the SNNs and full-replacement models. Responses of FC1 units to SF-filtered face images were examined at two different sizes (198 × 198, 99 × 99 pixels). (A) Six example FC1 units of SNNs with a different peak shift (PS). (B) Two-dimensional histograms of peak SFs at large images versus small images for the SNNs (left) and the full-replacement models (right). Solid lines indicate peak shifts of 0, and dashed lines indicate peak shifts of 1. (C) Distribution of peak shifts of units in the 20 SNNs (left) and the 20 full-replacement models (right). Arrowheads indicate the estimated locations of multiple peaks in the distribution (solid: major peaks, open: statistically detected but less obvious peaks). Gray columns indicate units with a response peak at SFs below 2 cycles/object for large and/or small stimuli.

We plotted the peak positions of 2,401 FC1 units of the 20 SNNs in a two-dimensional space defined by the peak SF for the large stimuli on the abscissa, and the peak SF for the small stimuli on the ordinate (Fig. 5B, left). Note that 46% of FC units were excluded from this analysis, either because they were not sensitive to SFs (23%) or because the largest responses were found at the end of the examined range of SFs and the peak SFs could not be determined (23%). The diagonal solid line in Fig. 5B represents the responses of a peak shift of 0, and the dashed line next to it represents the responses of a peak shift of 1. FC1 units of the SNNs were clustered in multiple groups in this scatter plot. One conspicuous group was selective to low SFs and was centered on the diagonal, i.e., peak shift values around 0. Another group was selective to higher SFs, and was clustered on the dashed line indicative of peak shift values around 1. The multimodality of the distribution can also be seen in the histogram (Fig. 5C, left). We applied an excess mass test for multimodality (Ameijeiras-Alonso et al., 2019, 2021) to this distribution. This test statistically determines the number of peaks in the distribution, with the null hypothesis that the true number of peaks is *N* (*N* = 1, 2, 3, …). The true number of peaks is estimated as the smallest *N* under which the null hypothesis is not rejected. The excess mass test also estimates the locations and heights of peaks from Gaussian kernel density estimation. The test revealed that there were three peaks in the distribution of the SNNs (first *p*-value < 0.001, second *p*-value < 0.001, third *p*-value = 0.096). Based on the probability density function derived from the histogram (Ameijeiras-Alonso et al., 2021), the peaks were estimated to be located at *−*4.85, 0.144, and 0.909 (open and solid arrowheads in Fig. 5C, left). Units sensitive to low SFs below 2 cycles/object were most frequent around a peak shift of 1 (gray columns). Comparing this result and the density map, the peak around 0 was mostly from the low spatial frequency group and the peak around 1 was from the high spatial frequency group. Although the third peak at the far periphery (at *−*4.85, open arrowhead) was statistically detected, it was much smaller in height than the other two peaks (1.1% of the peaks near 0 and 1). The results indicate that FC1 contained two major groups of units, those sensitive to low SFs, encoding SFs in the object-based coordinate, and those sensitive to high SFs, encoding SFs in the retina-based coordinate.

The distribution of peak shift values was drastically altered in the full-replacement models. In the two-dimensional plot shown in Fig. 5B (right), most data points were diffusely distributed in an elongated area between the diagonal and dashed lines, indicating that the SF reference frame of most units was intermediate between retina-based and object-based. An excess mass test again detected three peaks located at *−*4.84, 0.436, and 4.98 (Fig. 5C right, solid and open arrowheads; first *p*-value < 0.001, second *p*-value < 0.001, third *p*-value = 0.096). The second and third peaks at *−*4.84 and 4.98 (open arrowheads) were smaller than the primary peak at 0.436 (3.1% and 2.6% of the primary peak, respectively), making the distribution nearly unimodal.

Given the change of the peak shift distribution in the full-replacement models, we next analyzed one- and two-property replacement models to determine which subcortical properties were essential for the multimodal distribution of the SNNs. All these modified models exhibited unimodal distributions of the major peak (solid arrowheads) at different peak positions (Fig. 6). An excess mass test for multimodality detected two other less obvious peaks (open arrowheads) in each model as in the cases of the SNNs and the full-replacement models (Fig. 5C). The heights of these smaller peaks were 2.6–26 % of those of the major peaks, and were located at the periphery of the distribution.

**Figure 6.**
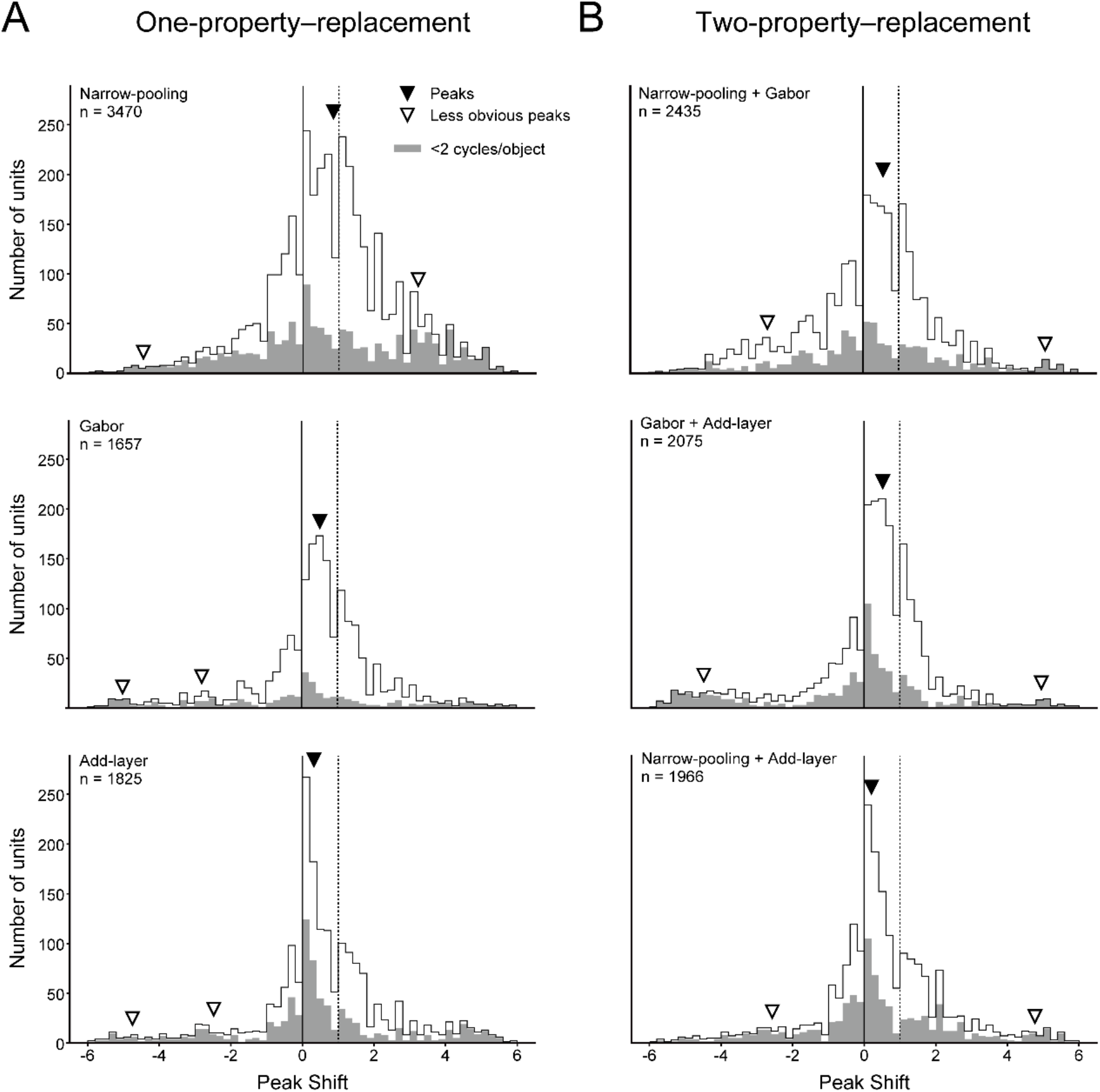
Distributions of peak shifts of FC1 units of the SNNs and one-property or two-property–replacement models. (A) Data from narrow-pooling model, add-layer model, and Gabor model. (B) Data from models with two modifications: narrow-pooling + Gabor, Gabor + add-layer, narrow-pooling + add-layer. Gray columns indicate units with a response peak at SFs below two cycles/object for large or small stimuli. Arrowheads indicate the estimated locations of multiple peaks in the distribution (solid: major peaks, open: statistically detected but small peaks). Gray columns indicate units with a response peak at SFs below 2 cycles/object for large and/or small stimuli.

Each of the three one-property–replacement models showed a characteristic distribution of the peak shift values. The narrow pooling models contained units with peak shift values between 0 and 1 in addition to units with peak shift values around either 0 or 1. The distribution became unimodal and broad, and was estimated to be centered at 0.85. In the Gabor models, units with peak shift values intermediate between 0 and 1 were the most abundant with a smaller number of units of peak shift values around 0 and 1. The distribution peak was estimated at 0.51. In the add-layer models, units with peak shift values around 0 were predominant, and exhibited a sharp distribution peak at 0.32. As to the two-property–replacement models, the narrow-pooling + Gabor models and the Gabor + add-layer models showed a broad distribution straddling the peak values from 0 to 1 (peak for the former, 0.56; peak for the latter, 0.52), whereas the narrow-pooling + add-layer models showed a sharp distribution peak at 0.20. As in the SNNs and the full-replacement models, units sensitive to low SFs (below 2 cycles/object) were most frequently found around the peak shift of 0 in all of the one- and two-property replacement models (gray columns). The results indicate that all of the three computational properties were responsible for the multimodal distribution of peak shift values observed in the SNNs. In particular, the smaller number of units with peak shift values around 1 in the Gabor models and the add-layer models suggests that the shallowness and the DoG-type filters were critical for preserving the unit sensitivities to retina-based SFs. The broad distribution observed for the narrow-pooling models and the narrow-pooling + Gabor models suggests that the wide pooling employed in the SNNs contributed to the two peaks at 0 and 1, by reducing units with peak shift values intermediate between 0 and 1.

### Effects of max pooling on SF tuning

We showed above that FC1 units of the SNNs were roughly grouped into two populations in terms of the reference frame of SF encoding. Because the max pooling yields the same output from a population of convolution layer units in response to slightly different spatial arrangement of local features, the max pooling operation is likely to affect the encoding of global configuration of face components. This information of global configuration will be reflected in a low range of SFs. Therefore, we next compared the effect of max pooling on the representation of SFs across different SF ranges.

We first analyzed the responses of the 96,800 units (32 filters × 55 × 55 resolution) in the first convolution layer. We obtained the response patterns across these units by feeding bandpass-filtered faces of two sizes (198 × 198 and 99 × 99 pixels; Fig. 2B) to the models, and quantified the difference between the SF tunings obtained for the two stimulus sizes by calculating the dissimilarity index (see Materials and Methods). In the SNNs with DoG filters, the dissimilarity index was high (around 0.6) for a low SF range up to approximately four cycles/object, but gradually decreased over a higher range of SFs (Fig. 7A, black curve). In the pooling layer, the dissimilarity index of the 6,272 units (32 filters × 14 × 14 resolution) became lower for a low SF range of less than four cycles/object than that of the convolution layer. For a high SF range of greater than four cycles/object, by contrast, it became higher than that of the convolution layer (Fig. 7A, compare the orange and cyan curves with the black curve). Thus, max pooling resulted in the SF tuning becoming similar between the two stimulus sizes for a low SF range, consistent with the results of peak shift analysis (Fig. 5C). Although these changes were observed both for wide pooling (5 × 5) and narrow pooling (3 × 3), the effects were larger for the former than for the latter (Fig. 7A, compare the orange curve with the cyan curve). This was more evident when we plotted the ratio of dissimilarity indices between before and after pooling (Fig. 7B). Furthermore, the ratio curve for wide pooling had smaller standard deviations than that for narrow pooling (shown as shades in Fig. 7B), indicating that wide pooling exerted its effects more consistently across the 15 individual faces and the seven facial expressions than narrow pooling.

**Figure 7.**
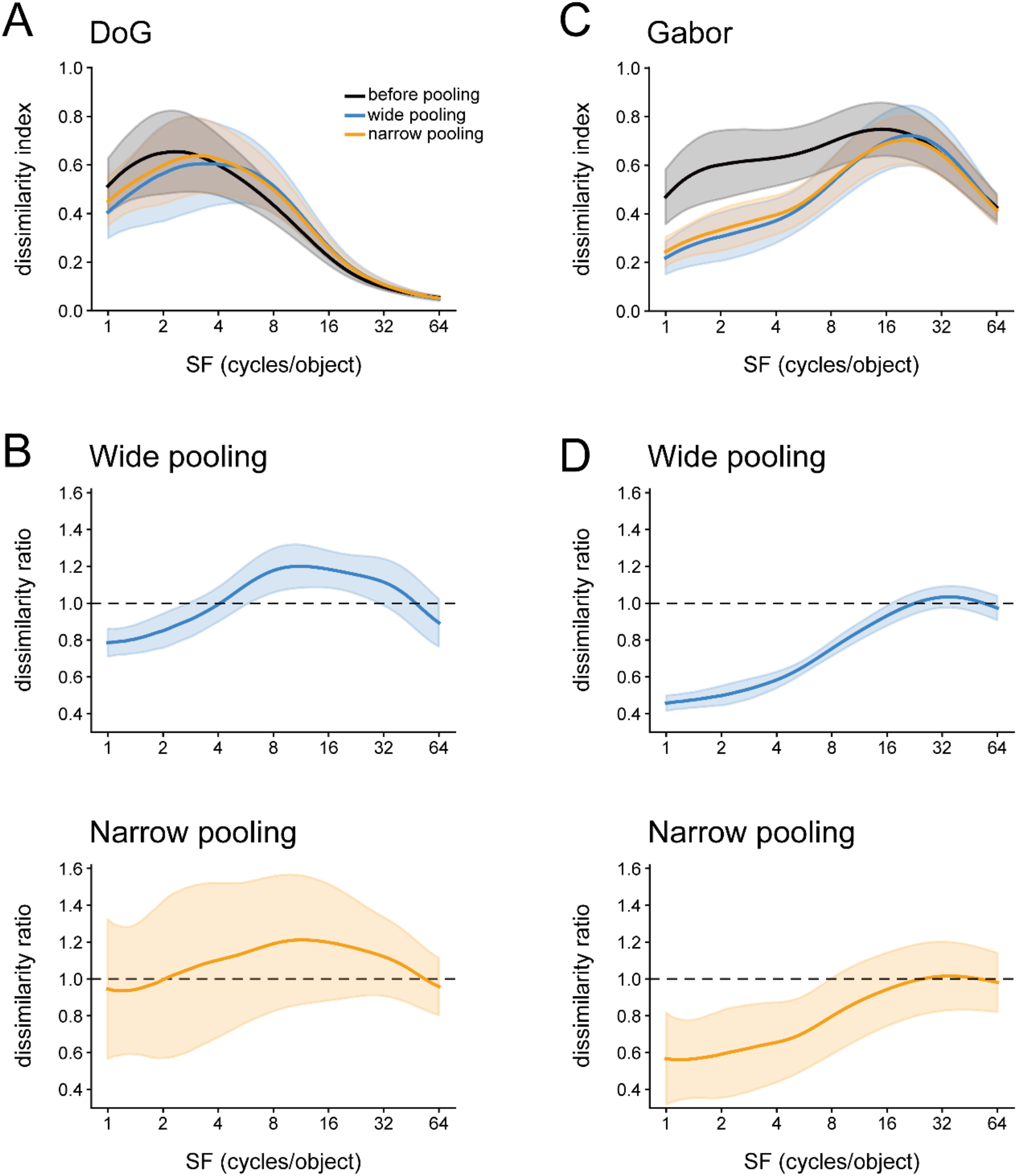
Effects of max pooling on the size-invariant responses to SFs. (A) Dissimilarity index curves of responses of units in the first convolution layer of the SNNs before max pooling operation (black), after wide pooling (blue), and after narrow pooling (orange). Dissimilarity indices, defined by the Euclidean distances of unit responses between different stimulus sizes (see Method and Methods), are plotted against the center SFs of input images. Solid lines indicate the means of dissimilarity indices across the seven facial expressions. Shades indicate standard deviations. Each dissimilarity index was normalized by the number of units and the maximum values. (B) Dissimilarity ratios of inputs and outputs of the max pooling operation (upper, after wide pooling; lower, after narrow pooling). (C, D) Data from units in the first convolution layer of the Gabor models. The conventions are the same as in A and B.

By contrast, the convolution-layer units of the Gabor models exhibited a constantly high dissimilarity index over most of the SF range (Fig. 7C, black). However, when we applied max pooling with windows of either 5 × 5 or 3 × 3 in size, the dissimilarity index became small over almost the entire SF range, with the largest decrease for 1–16 cycles/object (Fig. 7C; compare the orange and blue curves with the black curve). As in the case of the SNNs, the effect was stronger for wide pooling than for narrow pooling (Fig. 7D). These results demonstrated that max pooling rendered the SF tuning more invariant to stimulus size for units sensitive to low SFs, enabling them to represent SFs in the object-based coordinate. Regardless of the filter type in the first convolution layer (i.e., DoG vs. Gabor), wide pooling was more effective than narrow pooling in creating this response property.

### Effects of alternation of sliding strides on SF tuning

Finally, we examined the effect of another free parameter of our models, the stride size, on the SF sensitivity of FC1 units. We changed the stride of the two max pooling layers of the SNNs from 4 to 2. The stride size of 2 was also employed in the narrow-pooling model. This modified model with a smaller stride of 2 achieved a mean correct rate of 0.54, which was better than the SNNs (0.51) but similar to the narrow pooling models (0.55) (vs. SNN, *p* = 0.0020; vs. the narrow pooling, *p* = 0.23; *t*-test with Bonferroni correction).

An analysis of FC1 responses to two stimulus sizes (198 × 198 and 99 × 99 pixels) revealed that the distribution of the peak shift had three peaks, at *−*4.17, *−*0.0107, and 0.976 (first *p*-value < 0.001, second *p*-value < 0.001, third *p*-value = 0.098; excess mass test for multimodality; solid and open arrowheads in Fig. 8). Two of them were conspicuous and located near 0 or 1 (solid arrowheads), and the third one at the periphery of *−*4.17 (open arrowhead) was small (3.2% and 3.3% of the two major peaks). Comparisons with Fig. 5B, C show that the small-stride model exhibited a similar SF representation as in the SNNs, in that there were two main groups of units, one sensitive to low SFs, representing SFs in the object-based coordinate (peak shift around 0), and the other sensitive to high SFs, representing SFs in the retina-based coordinate (peak shift around 1). The change of the stride from 4 to 2 had little effects on the reference frame of SF sensitivity of FC1 units.

**Figure 8.**
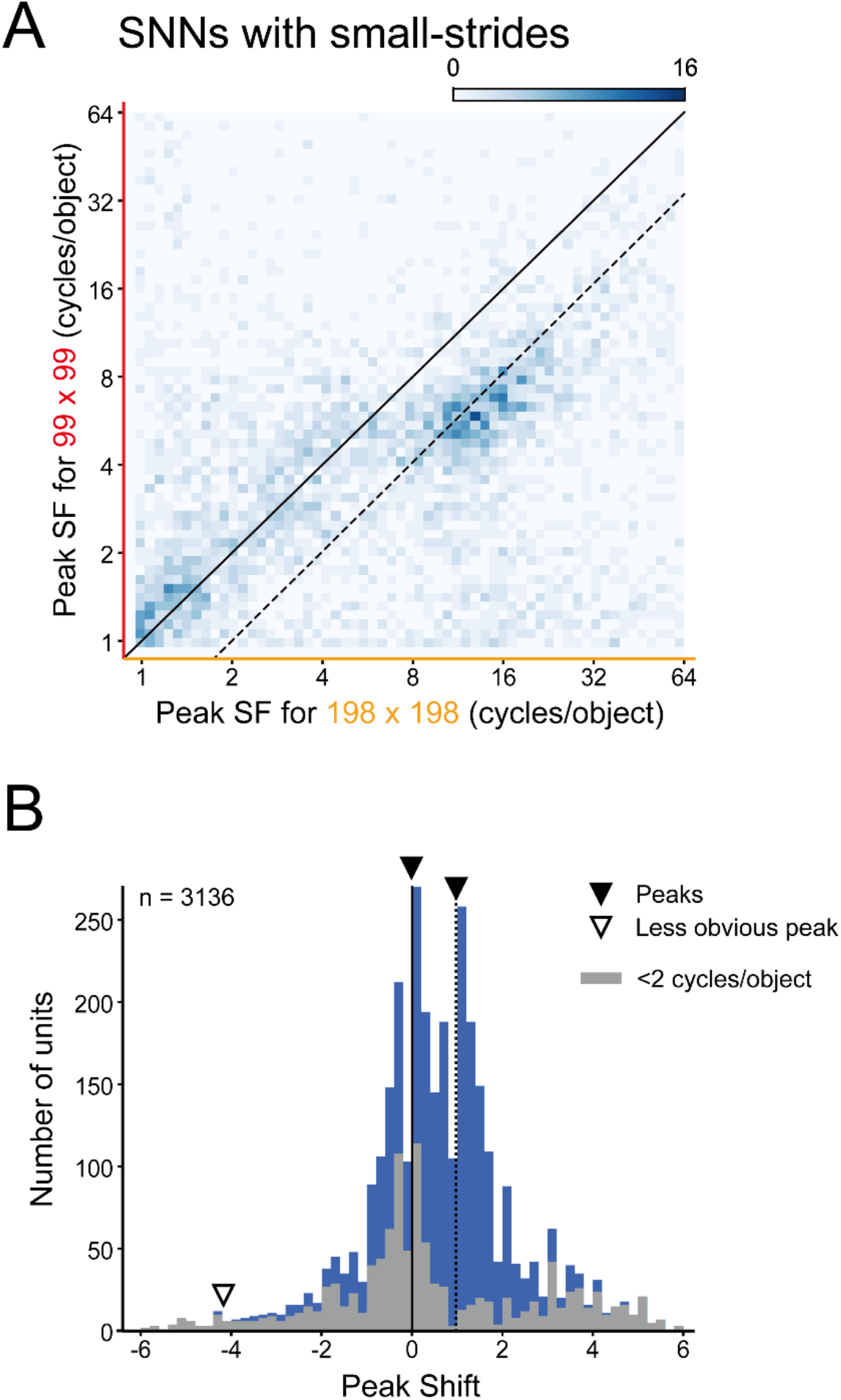
Effects of alternation of sliding strides of pooling windows on SF tuning reference frames of FC1 units of the SNNs. Responses of FC1 units were obtained in the same way as in Fig. 5. (A) A two-dimensional histogram of peak SFs obtained with large (198 × 198) versus small (99 × 99) stimulus images. (B) Distribution of peak shifts of units. Arrowheads indicate the estimated locations of multiple peaks in the distribution.

## Discussion

We analyzed the ability of the SNNs and modified models to classify facial expressions, with the goal of determining what architectural or physiological properties underlie the modest performance of facial expression discrimination supported by the primate subcortical pathway. The SNNs were implemented with the three prominent subcortical properties, i.e., shallow processing, DoG-type filters at the first processing stage, and spatial pooling over wide areas (Fig. 1). The SNNs classified the seven basic facial expressions with modest performance (Fig. 3). Replacement of any one of these properties with the corresponding cortical properties resulted in higher performances (Fig. 4). Replacement of a combination of two or three properties further improved classification performances in a partially additive manner (Fig. 4). These results suggest that all three subcortical properties of the SNNs underlie the modest performance. A major group of units in the final processing layer (FC1) of the SNNs was sensitive to SFs defined in the retina-based coordinate, whereas another group responding to low SFs encoded SFs in the object-based coordinate (Fig. 5). The number of retina-based units was reduced in most of the modified models, suggesting that the three features are also important for preserving retina-based SF information (Figs. 5, 6). Max pooling over the wide window employed in the SNNs contributed to object-based SF representation of units sensitive to low SFs (Fig. 7). These findings advance our understanding of the computational processes utilized by the subcortical pathway in facial expression recognition.

### Modest performance of the SNNs and neural computations of the subcortical pathway

Based on psychological assessment and brain imaging in V1-lesioned patients, it has been proposed that affective blindsight is mediated by components of the subcortical pathway spared by the lesions, including the superior colliculus, pulvinar, and amygdala (de Gelder et al.,1999; Pegna et al., 2005; Striemer et al., 2019). One view assumes that the shortest route directly connecting the three subcortical structures conveys facial expression information from the superior colliculus via the pulvinar to the amygdala (Tamietto and de Gelder, 2010). A different view proposes that information from the pulvinar reaches the amygdala through the facial processing system in the temporal cortex, under an assumption that “the direct connections of the pulvinar with the amygdala are likely insufficient in themselves for recognizing emotional expressions” (Gerbella et al., 2019). The present study demonstrates that the SNNs with only three processing stages and the subcortical physiological properties can successfully acquire an ability to discriminate facial expressions.

The average correct rate of classifying the seven facial expressions in the present study was 0.51. This rate was well above chance (1/7 = 0.14), but was far from perfect. The modest correct rate is in line with the performance of patients with affective blindsight. Pegna et al. (2005) reported that a patient with bilateral lesions in V1 discriminated happy faces from either angry, sad, or horrified faces at correct rates of 0.58–0.62, marginally above the chance level of 0.5. Another patient with bilateral lesions in V1 exhibited correct rates of 0.64–0.67 for happy vs. fearful or angry faces (chance level = 0.5; Striemer et al. 2019). The residual ability of facial expression classification in these patients was only moderate compared to the nearly perfect performance in healthy people. This raises the question of why subcortical processing supports vision more poorly than visual functions mediated by the cortical pathway.

A traditional explanation is that neurons in the subcortical pathway respond to low SFs and are less sensitive to high SFs than the cortical pathway (e.g., Vuilleumier et al., 2003; Méndez-Bértolo et al., 2016; Burra et al., 2019). This will limit the ability of the subcortical pathway to analyze the fine details of visual images, and can itself result in the inaccurate processing of face images. However, the dependence of the subcortical response on low SFs has been disputed by other researchers (De Cesarei and Codispoti, 2013; McFadyen et al., 2017). Our results suggest that low SF sensitivity, if important, was not the only cause, because the DoG filter models combined with narrow-pooling or add-layer modifications exhibited improved performances, despite the fact that our DoG filters were tuned to low SFs, and had full width at half maximum of 0.067–1.0 cycles/degree. Note that we estimated this value on an assumption of the image size of 30.5° based on our DoG parameters, the filter resolution, and the RF size of superior colliculus neurons representing the foveal region. The range of DoG-filter width corresponds to that applied in models of the superior colliculus in a recent simulation study (Méndez et al., 2022).

Another explanation is that the small number of processing stages in the cortical pathway hampers detailed analysis of visual inputs. However, a previous study (Dailey et al., 2002) showed that CNNs that had only two processing layers, with Gabor filters at the first stage, performed highly accurate discrimination of facial expressions (the mean correct rate for classifying six facial expressions was 0.90). The performance of our SNNs incorporating the three subcortical properties was not this high. This was not due to inadequate training, because the performance reached a plateau and stayed stable over a large number of iterations in the training sessions (Fig. 3). This was further verified by showing that the correct performance for the test set remained unchanged even after overly excessive training with 3,000,000 iterations. Furthermore, replacing not only the small number of processing layers but also the filter type at the first processing layer and the width of the pooling window with the corresponding cortical properties improved the performance of the SNNs (Fig. 4). The three properties at least partially underlie the less accurate processing of facial images in the subcortical pathway, and hence, may be responsible for the low performance in affective blindsight.

### Confusions of facial expressions in the SNNs, DNNs and patients

The classification accuracy of the SNNs varied across facial expressions (Fig. 3A, B). The classification performance of the SNNs was best for happy and surprised faces and worst for sad and neutral faces. The rank order of performance on the seven facial expressions was largely consistent across the 20 SNNs trained independently from random states (Fig. 3C). It also corresponds to the classification performance by previously developed AlexNet-based DNNs (Inagaki et al., 2022b). These DNNs were trained to discriminate between the seven expressions derived either from the KDEF database or the Kokoro Research Center (KRC) facial expression database (Ueda et al., 2019). Like the SNNs, the DNNs exhibited the best performance for happy and surprised faces (KDEF: 0.93 for happy, 0.84 for surprised; KRC: 0.91 for happy, 0.84 for surprised; chance level, 0.14). This coincidence may simply suggest that within each database, facial features are consistent across faces with happy or surprised expressions, but are more diverse across faces with sad or neutral expressions. However, the variations across examples of facial expressions within a database are not the sole reason for the difference in the performance across facial expressions, because neutral faces were classified poorly by the SNNs (Fig. 3B; correct rate = 0.34), but the DNNs of Inagaki et al. (2022b) classified them with high correct rates (0.85 for KDEF, 0.82 for KRC). An alternative, yet-to-be-tested explanation is that the ease (or difficulty) of classification may vary across the facial expressions owing to differences in the conspicuousness of component facial actions underlying various expressions. Similarities between neural networks regarding expression-specific performance may vary according to these differences.

The relatively poor ability to distinguish between sad and neutral faces was also observed in another CNN with the first layer of DoG filtering and average pooling (Méndez et al., 2022). This CNN was constructed to simulate facial processing in the superior colliculus, and was trained to discriminate three facial expressions: happy, sad, and neutral. The CNN showed the best performance for happy faces and moderate performance for sad faces, but classified neutral faces into neutral faces with a classification rate of 0.49 and into sad faces with a rate of 0.39. The fact that this CNN and the SNNs in the present study demonstrated this confusion, whereas AlexNet-based DNNs and our add-layer models (0.52 for sad, 0.67 for neutral) did not, suggests that the convolution processes after the initial DoG filtering (in the case of add-layer models) or the convolution by the Gabor filters (in the case of AlexNet-based DNNs) may be critical for classification of sad and neutral faces.

Finally, we point out that the expression-dependent performance of the SNNs also had both similarities and dissimilarities to that observed in a V1-lesioned patient. The patient reported by de Gelder et al. (1999) classified happy and sad faces with a higher correct rate than angry and fearful faces; our SNNs and this patient classified happy faces well, whereas the performance for sad faces was poor in the SNNs but good in the patient.

### Reference frame of coding SF information and invariance of visual responses

FC1 units of the SNNs consisted of two major groups, each with different properties regarding SF processing (Fig. 5). One group of units responded best to the same object-based SFs (cycles/object) regardless of the stimulus size (peak shift around 0). This size-invariant response indicates that these units represent SFs in the object-based coordinate. Most of these units were tuned to a low SF range (around one to two cycles/object). In the other group of units, the optimal object-based SFs shifted when testing was performed with different stimulus sizes. The direction of the shift was consistent with the interpretation that the units were tuned to retina-based SFs (cycles/degree) (peak shift around 1). That is, for larger stimuli, the units responded to higher object-based SFs that corresponded to the same retina-based SFs. The DoG filters at the initial stage and the shallow architecture appear to be critical for preserving the SF representation based on the retina-based coordinate, because FC1 units with retina-based SF sensitivity were reduced in number when the first convolution layer were changed to Gabor filters or when the number of processing layers was increased (Fig. 6A, middle, bottom).

A major group of the object-based SF units in the SNNs were tuned to low SFs (Fig. 5B). This curious bias of the object-based units towards low SF sensitivity likely resulted from the wide max pooling process. Lowpass-filtered facial images contain only coarse structure such as solid blobs at eye or mouth positions. Positional information of these blobs is initially detected by DoG filters, and is encoded as response patterns across units in the convolution layer. These blobs appear in different positions and scales for images of different sizes, and thus the response patterns vary between different sizes. After the max pooling operation, however, response patterns would become more similar between different sizes, because this operation renders units in the pooling layer insensitive to slight changes in spatial arrangement of local features. Indeed, the dissimilarity index for lowpass-filtered images decreased after max pooling in our data (Fig. 7). This effect might result in object-based SF tuning (i.e., preferential responses invariant of image size to a particular range of object-based SFs) for lowpass-filtered images. Wider pooling window would enhance this effect at the expense of losing fine details of inputs. When the pooling window is narrower, this effect would be incomplete, and units with intermediate peak shift values would increase, as we found in the narrow-pooling models (Fig. 6A, top).

One may wonder why FC1 units of the SNNs maintained sensitivity to retina-based SFs, i.e., size-dependent representation of SFs, despite the demand that we imposed on the SNNs to classify facial expressions regardless of the seven different face image sizes. One possibility is that the architecture of our SNNs cannot achieve sufficient object-based representation, and remains suboptimal for the required task even after the excessive training sessions. This may be a reason for the modest classification performance of the SNNs. Indeed, replacement of the subcortical processing properties with the cortical properties resulted in the representation becoming more object-based (Fig. 5B, C, Fig. 6) and improved the classification performance (Fig. 4). However, if object-based SF encoding was the only requirement for optimal performance under our training conditions, the models that showed object-based SF encoding should have had the highest correct rate, but this was not the case. The add-layer models and the narrow-pooling + add-layer models exhibited the best object-based encoding of SFs (Fig. 6A bottom, 6B bottom), while they performed worse than the full-replacement model (Fig. 4A). The representation acquired for the classification depended not only on the task demand of size-invariant classification of facial expressions, but also on other, yet unspecified, constraints deriving probably from the architecture of the models.

In the primate amygdala, the responses of many neurons are affected by retina-based SFs, and only a minority of neurons have perfect object-based SF sensitivity (Inagaki and Fujita, 2011). By contrast, many FC1 units tuned to low SFs of the SNNs exhibited object-based SF sensitivity. The paucity of evidence for units with object-based SF sensitivity in the amygdala may be related to the fact that the previous electrophysiological study (Inagaki and Fujita, 2011) did not present face images with very low SFs, and may have overlooked the neurons with object-based SF sensitivity in this range of SFs.

Some inferior temporal cortex neurons exhibit invariant responses to changes in shape sizes (Rolls and Baylis, 1986; Ito et al., 1995). The max pooling operation may help achieve these invariant responses. To some degree, max pooling ignores positional changes of inputs in each region of interest. Because size changes involve alternations in edge positions without modifications in topologies, if the changes are small enough to be covered by each region of interest, stimuli before and after the changes would yield similar responses. The effects of wide pooling shown in Figure 7 suggest that some aspects of the invariant responses of inferior temporal cortex neurons can simply be achieved by bypassing early cortical areas with high spatial resolutions such as V1. Such shortcut routes indeed exist, including the projection from the pulvinar to V2 and then to the posterior inferior temporal cortex and the projection from the pulvinar to V4 and then to the anterior inferior temporal cortex (Pessoa and Adolphs, 2010).

The size invariance in low SFs is important in newborns. They have blurred visions that relies on low SFs (Atkinson et al., 1974; Dobson & Teller, 1978), but respond to faces or face-like patterns irrespective of the stimulus size or the viewing distance (Cassia et al., 2001; De Heering et al., 2008). These findings indicate that the ability of size-invariant face recognition based on low SFs is innately implemented in our visual system. Convergence of inputs from the superficial layer of the superior colliculus to the deep layer, which is already present in newborns (Wallace et al., 1997), may be part of the neural substrate supporting this aspect of size-invariant face recognition.

### Concluding remarks

The present study provides the first computational model for facial expression processing along the subcortical pathway (see Méndez et al., 2022 for a model of face processing in the superior colliculus). Despite the celebrated success of DNNs in modeling visual processing in the ventral cortical pathway, it has remained unclear whether and how the CNN architecture can be adapted to processing in the subcortical pathway. We demonstrated that the SNNs implemented with the three computational properties of the subcortical pathway, i.e., a shallow layer architecture, concentric receptive fields at the first processing stage, and a greater degree of spatial pooling, were successfully trained to discriminate facial expressions with a modest correct rate. The three properties were all essential for reproducing the modest performance seen in V1-lesioned patients, as well as the representation of SFs in the retina-based coordinate observed in a population of amygdala neurons. Research interest in the role of subcortical structures in cognitive functions has recently surged, but physiological data are still much sparser for subcortical structures than for the cerebral cortex (Janacsek et al., 2022). Computational approaches such as the one we present here are expected to partially compensate for this data scarcity and to guide future research.

## Acknowledgments

This work was supported by grants from the Ministry of Education, Culture, Sports, Science and Technology of Japan (JP17H01381 and JP21H02596 to IF; JP18H04197, JP20H04578, and 20K12023 to MI); the Center for Information and Neural Networks; the Ministry of Internal Affairs and Communications of Japan. CL was supported by the Research Fellowship for Young Scientists from the Japan Society for the Promotion of Science.

## Author contributions

CL, MI, TS, and IF designed the research; CL, MI, and TS performed the research; CL, MI, TS, and IF wrote the paper. All authors approved the submitted version.

## Data availability

All data and analysis codes are available from the corresponding author upon request.

## Ethics statement

Written informed consent was obtained for the publication of any identifiable images included in this article.

